# Systematic evaluation of anatomical details on transcranial electric stimulation and transcranial magnetic stimulation induced electric fields in a non-human primate model

**DOI:** 10.1101/2022.12.05.519206

**Authors:** Kathleen E. Mantell, Sina Shirinpour, Oula Puonti, Ting Xu, Jan Zimmermann, Sarah R. Heilbronner, Axel Thielscher, Alexander Opitz

**Affiliations:** Department of Biomedical Engineering, University of Minnesota, Minneapolis, USA; Danish Research Centre for Magnetic Resonance, Centre for Functional and Diagnostic Imaging and Research, Copenhagen University Hospital - Amager and Hvidovre, Copenhagen, Denmark; Center for the Developing Brain, Child Mind Institute, New York, USA; Department of Neuroscience, University of Minnesota, Minneapolis, USA; Department of Health Technology, Technical University of Denmark, Kgs. Lyngby, Denmark

**Keywords:** Noninvasive brain stimulation (NIBS), Transcranial Magnetic Stimulation (TMS), Transcranial Electric Stimulation (TES), Non-human Primate, Finite Element Method Modeling, Ultra-high Field Imaging

## Abstract

Non-human primates (NHPs) have become key for translational research in noninvasive brain stimulation (NIBS). However, in order to create comparable stimulation conditions to humans it is vital to evaluate and match electric fields across species. Numerical models to simulate electric fields are an important tool for experimental planning in NHPs and translation to human studies. It is thus essential to evaluate how anatomical details in NHP models affect NIBS electric fields. Here, we create a highly accurate head model of a non-human primate (NHP) from 10.5 T MR data. We systematically evaluate how muscle tissue and head size (due to different MRI fields of view) affect simulation results in transcranial electric and magnetic stimulation (TES and TMS). Our findings indicate that the inclusion of muscle can affect TES electric field strength up to 29.5% while TMS is largely unaffected. Additionally, comparing a full head model to a cropped head model illustrates the impact of head size on electric fields for both TES and TMS. We find opposing effects between TES and TMS with an increase up to 13.5% for TES and a decrease up to 21.5% for TMS for the cropped head model compared to the full head model. Our results provide important insights into the level of anatomical detail needed for NHP head models and can inform future translational efforts for NIBS studies.

**Highlights:**

- We created a high resolution non-human primate FEM head model from 10.5 T MR data.
- We ran transcranial electric and magnetic stimulation (TES and TMS) simulations to evaluate the effects of muscle and head size on the induced electric field in the brain.
- We simulated both isotropic and anisotropic muscle conductivities.
- Muscle tissue can greatly affect simulation results for TES (increase electric field strength by up to 29.5%), but not for TMS.
- There are opposing, but large effects of using a full head model versus a cropped head model for both TES (increase up to 13.5%) and TMS (decrease of up to 21.5%)

## Introduction

Transcranial electric stimulation (TES) and transcranial magnetic stimulation (TMS) are two popular noninvasive brain stimulation (NIBS) techniques to modulate and study brain activity in humans. TES uses electrodes placed on the scalp to apply an electric current which creates an electric field in the brain (Opitz et al., 2015). TES electric fields can modulate ongoing brain activity and bias spike timing (Johnson et al., 2020; Krause et al., 2019). TMS uses a magnetic coil placed on the scalp to generate a time varying magnetic field that induces an electric field in the brain (Barker et al., 1985; Opitz et al., 2011). TMS electric fields can be at suprathreshold levels that can directly cause neurons to fire action potentials (Di Lazzaro & Rothwell, 2014; Mueller et al., 2014).

NIBS methods are used in a wide range of clinical applications and cognitive neuroscience. However, despite their increasing use, NIBS physiological mechanisms are still incompletely understood. Research in non-human primates (NHP) models has strongly advanced our understanding of NIBS physiology (Johnson et al., 2020; Kar et al., 2017; Krause et al., 2017, 2019; Mueller et al., 2014; Romero et al., 2019). Invasive recordings in NHPs enable researchers to measure NIBS effects on the level of single-units and local field potentials typically not possible in humans. However, in order to translate findings from NHPs to humans, it is crucial that electric fields are comparable across species. Due to differences in head anatomy, electric fields will differ between NHPs and humans, requiring careful consideration for experimental planning (Alekseichuk et al., 2019).

Computational modeling has become one of the most prominent ways to study NIBS induced electric fields. From MRI segmentation masks, anatomically realistic 3D models are created that can distinguish between several head and brain tissues and their respective conductivities. Numerical methods such as the finite element method (FEM) allow accurate simulations of NIBS electric fields for specific stimulation configurations and head models. Ultra-high field MRI scans enable imaging of NHP anatomy in unprecedented detail (Grier et al., 2022). In the present work, we leverage 10.5T anatomical MRI scans to create a detailed NHP head model.

In previous work, we investigated the effects of overall brain volume and head size on NIBS electric fields (Alekseichuk et al., 2019). However, we did not investigate the effect of anatomical structures such as muscle tissue and muscle anisotropy. NHPs have a large volume of muscle around the skull, which can affect electric fields delivered to the brain in particular for TES. The conductivity of tissues between electrodes plays a large role in how much current shunting occurs (Opitz et al., 2015; Santos et al., 2016; Saturnino et al., 2019), thus the ratio of skin to muscle conductivity is an important interface to investigate. Additionally, muscles are fibrous tissues with the fibers aligned in specific directions. This results in anisotropic muscle conductivity (Gabriel et al., 2009). Here we evaluate the effect of modeling muscle tissues with and without anisotropic conductivities. We expect that TES simulations will be affected by the presence of muscle tissue as it will decrease the volume of the higher conducting skin layer and thus decrease current shunting. TMS, on the other hand, is not expected to have large changes because the magnetic field will be unaffected by the non-brain tissue conductivities.

An additional important factor for NHP head modeling is the field-of-view of the MRI and the inclusion of tissues such as neck and jaw in the FEM model. Many MR imaging protocols are optimized to only include brain tissues thus resulting in cropped head field of views. Cropped head images are also preferred for image registration across multiple animals and for training of automated segmentation pipelines due to increased variability outside of the brain region for NHPs. Additionally, it is significantly more time consuming to hand segment a full head MRI than a cropped head MRI. It is thus important to systematically evaluate how the NIBS generated electric fields are affected by the difference in overall head size. Based on our previous cross-species work (Alekseichuk et al., 2019), we expect to see a decrease in the induced electric field for a full head model with TES, but an increase for TMS.

In summary, the main goal of this study is to understand how the anatomical details of a NHP model affect NIBS electric fields. To this end we develop a first-of-its-kind, 10.5 T MRI-based 3D NHP model. We evaluate the effect of 1) including muscle tissue and muscle anisotropy and 2) modeling the whole head vs. a cropped head NHP model. This will help researchers to determine the amount of anatomical detail needed for precise electric field modeling in NHPs and inform experimental planning for future experiments.

## Methods

### Animal preparation

Experimental procedures were carried out in accordance with the University of Minnesota Institutional Animal Care and Use Committee and the National Institute of Health standards for the care and use of NHPs. All subjects were fed *ad libitum* and pair-housed within a light and temperature-controlled colony room. The subject was not water restricted and had no prior implant or cranial surgery. The subject was fasted for 14–16 hr prior to imaging. On scanning days, anesthesia was first induced by intramuscular injection of atropine (0.5 mg/kg), ketamine hydrochloride (7.5 mg/kg), and dexmedetomidine (13 μg/kg). The animal was transported to the scanner anteroom and intubated using an endotracheal tube. Initial anesthesia was maintained using 1.0%–2% isoflurane mixed with oxygen (1 L/m during intubation and 2 L/m during scanning to compensate for the 12-m length of the tubing used).

### MR Image and CT Scan Parameters

Structural imaging data was acquired for an anesthetized 4-year-old female *Macaca fascicularis*. The T1w structural scan was completed on a Siemens 10.5 T scanner with a 3D-MPRAGE sequence; FOV: 131 × 150 mm^2^; matrix size of 280 × 320 (0.5 mm isotropic resolution); TR/TE of 3300/3.56 ms (Grier et al., 2022). The CT scan was completed on a Siemens Biograph64. Axial slices had a voxel size of 0.34 × 0.34 mm^2^ and a resolution of 0.60 mm in the Z-direction. The matrix size was 512 × 512 × 222.

### NHP Segmentation and Head Model Generation

We started the segmentation process by first modifying the MR image using intensity normalization with Freesurfer’s mri_nu_correct.mni (Sled et al., 1998). The MR image and CT scan were registered using FLIRT (Jenkinson et al., 2002) and analyzed using various tools such as thresholding and other image processing routines to get initial baseline segmentation masks. We then completed the segmentation process with hand corrections in ITK-SNAP (Yushkevich et al., 2006). These masks were then upsampled to 0.25 mm^3^ voxels and smoothed using a gaussian kernel in python to remove small bumps resulting from inconsistencies in the hand segmentation process. Finally, we used the SimNIBS v4.0b1 meshmesh pipeline (Puonti et al., 2020) to generate a 3D model from the smoothed masks.

The muscles were segmented by hand. We elected to include muscles that are expected to be affected by typical TMS and TES protocols (Figure 1A). This included the occipitofrontal muscle, temporalis muscles, masseter muscles, and posterior muscles of the neck. Additionally, the lateral and medial pterygoid muscles were segmented for additional context for the temporalis and masseter muscles. These muscles were labeled based on both macaque and human muscle anatomy (Gilroy et al., 2012; Richmond et al., 2001; Schwartz & Huelke, 1963; Susan Standring, 2020). Facial muscles and most anterior muscles of the neck were not segmented.

**Figure 1.**
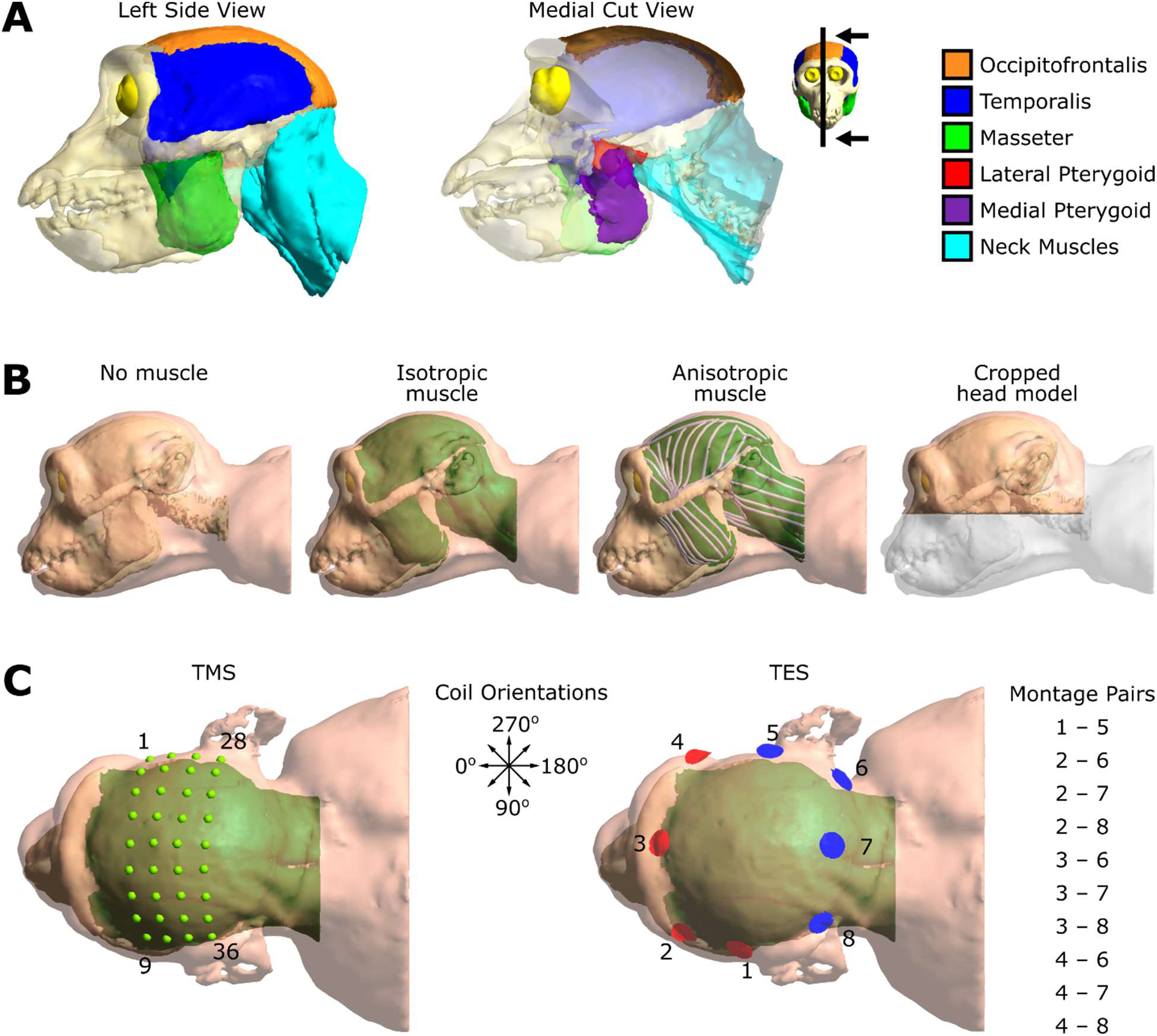
3D head model and simulation setup. A) Six classes of muscles were selected during the segmentation process (occipitofrontalis, temporalis, masseter, lateral pterygoid, medial pterygoid, and dorsal neck muscles). The medial cut view shows the lateral and medial pterygoid muscles; the inset figure shows the cropped plane and arrows indicate the viewing direction. The primary focus was the muscle that is directly under the TMS coil and TES electrode locations. B) Four variations of the 3D model were generated to compare muscle conditions (no muscle, isotropic muscle, and anisotropic muscle) and full (no muscle) vs. cropped models. No muscle is equivalent to only having skin outside the skull. Isotropic muscle has an isotropic muscle conductivity (0.160 S/m) for all muscle elements. Anisotropic muscle has an individualized conductivity tensor at each element (see tensor streamlines to visualize the directions of the largest eigenvectors). The cropped head model was generated from the same masks as the full head model, however, the masks were cropped in the superior/inferior and anterior/posterior axes. WM, GM, and CSF were kept intact except for a small section of the posterior region of the spinal cord. C) Simulation setup for TMS and TES. Left, TMS coil positions (green dots) were arranged in a 4×9 grid with 1 cm spacing using 8 orientations (black arrows, 0° to 315° in 45° increments). Right, TES anode and cathode electrodes were placed around the head and 10 different montages were selected to have 1 mA current pass through different volumes of muscle.

Muscle conductivity anisotropy is not a standard functionality in the current release of SimNIBS. However, SimNIBS is able to run simulations with anisotropic conductivities, so the source code was modified to allow muscle tissue to be anisotropic. We defined a reference muscle fiber in the muscle coordinate system with conductivity defined as follows (the fiber runs parallel to the x-axis):

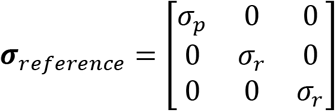

Where *σ_p_* is the conductivity parallel to the muscle fiber, and *σ_r_* is the conductivity radial to the muscle fiber. The isotropic muscle conductivity (σ_iso_) is 0.160 S/m (Gabriel et al., 2009). We opted to use a ratio of 5:1 (parallel : radial) conductivity based on data in (Gabriel et al., 2009). To ensure the mean conductivity was the same in the anisotropic simulations we used a volume normalized approach such that 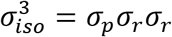, where *σ_p_* = 5*σ_r_*. Thus, σ_p_ = 0.4678 S/m and σ_r_ = 0.0936 S/m. We wrote custom MATLAB (MathWorks, Natick, MA) scripts to assign anisotropic conductivity tensors to each muscle triangle and tetrahedron, and isotropic conductivities to all other triangles and tetrahedrons for this specific NHP model. Muscle fibers were connected between each muscle’s origin and insertion areas. Based on this, the direction of the local muscle fiber was assigned to each triangle and tetrahedron as a vector. We used this muscle orientation vector to generate a rotation matrix from the reference muscle fiber to the assigned orientation. The rotation matrices were then used to calculate the conductivity tensor matrix for each element: 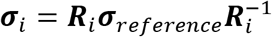, where **R**_i_ is the rotation matrix for the i^th^ element. Example tensor streamlines indicating the direction of the largest eigenvectors in the surrounding areas can be seen in Figure 1B, anisotropic muscle.

To study the effect of limited MRI field of view, we generated a cropped head model from the same smoothed masks as the full head model, however the mask data was removed beyond selected transverse and frontal planes (see supplementary Figure S1 to see the full and cropped masks overlayed on the MRI). The cropped mask was then input into the meshmesh pipeline to generate the cropped head model (see Figure 1B, cropped head model). A small portion of the WM and CSF in the spinal cord was lost in the crop, but no GM was cropped out. A comparison of volumes between the full and cropped head models can be found in Table 1.

**Table 1.** Full vs. cropped volume comparison. Cropping the head model resulted in a large reduction of volume, but most volume lost was limited to skin and skull. GM volume was unaffected by the crop, but a small portion of the caudal spinal cord was removed, resulting in a drop in WM and CSF volume in the cropped model.

| Model | Head Volume (cm<sup>3</sup>) | Brain Vol (WM+GM+CSF) (cm<sup>3</sup>) | GM Vol (cm<sup>3</sup>) | WM Vol (cm<sup>3</sup>) |
| --- | --- | --- | --- | --- |
| Full | 599.91 | 67.59 | 32.53 | 20.81 |
| Cropped | 244.42 | 66.86 | 32.53 | 20.25 |

### FEM Simulations

We ran all simulations using SimNIBS v3.2.3. Simulations were run to assess the differences in TMS and TES generated electric field estimates between the cases of no muscle, isotropic muscle, and anisotropic muscle, as well as between full head models and cropped head models. Simulations were performed using conductivity values for each tissue type as follows: σ_skin_ = 0.465 S/m, σ_muscle_ = 0.160 S/m, σ_bone_ = 0.01 S/m, σ_CSF_ = 1.654 S/m, σ_GM_ = 0.275 S/m, and σ_WM_ = 0.126 S/m (Gabriel et al., 2009; Windhoff et al., 2013). We used one full head model and its associated cropped head model for all simulations. In the case of no muscle, we changed the conductivity of the muscle tissue to that of skin. In the isotropic muscle case, we used 0.160 S/m for the conductivity of the muscle tissue. For anisotropic muscle cases, we customized the conductivity for each muscle element, as described above.

For TMS (Figure 1C, TMS), we generated a grid of 4×9 positions with 10 mm spacing which was placed across the top of the head. The grid spanned thick and thin muscle regions (temporalis and occipitofrontal, respectively). There were also 8 different coil orientations for each position, from 0° to 315° (where 0° is parallel to the to the longitudinal fissure, rotated counterclockwise in 45° increments). We used the Magstim 70 mm figure-of-eight coil included with the SimNIBS package with a dI/dt = 1 × 10^6^ A/s to simulate the electric fields.

For TES (Figure 1C, TES), 8 electrode locations were selected around the head, 4 of which were anodes, and 4 cathodes. Electrodes were set to be +/- 1 mA and were 1cm diameter, 5 mm thick simple circle electrodes. We selected anode cathode pairs such that a current passing through the two electrodes would go through different amounts of muscle tissue at different angles. In total we selected 10 pairs, 1–5, 2–6, 2–7, 2–8, 3–6, 3–7, 3–8, 4–6, 4–7, and 4–8.

### Data Analysis

Electric field simulation results were analyzed using custom MATLAB scripts. During the analysis, we excluded three TMS grid positions (1, 28, and 36) that were not feasible due to the coil intersecting the head model.

To compare the simulation results between two different simulations, we subtracted the results of one simulation from the results of the other at each GM element. The percent change was then calculated by dividing each of those difference values by the electric field strength robust maximum value of the standard condition (e.g., no muscle condition or cropped head model).

We defined the robust maximum of the electric field strength (|E|_max_) to be the 99.9^th^ percentile value of the electric field strength in all GM tetrahedra for a given simulation. This is done to ensure that the maximum is not driven by a numerical inaccuracy in the FEM simulation.

Since the same head model was used for all three muscle conditions, elementwise data could be compared one-to-one between models. We created the cropped head model from the same MRIs and cropped versions of the segmentation masks; however, it did not have the same triangle and tetrahedron configuration due to the rerun of meshmesh. Thus, simulation results had to be interpolated at the locations of the gray matter triangle and tetrahedron elements of the full head model in order to do one-to-one comparisons. We used the SimNIBS get_fields_at_coordinates function to interpolate the results into the full head GM volume, where the centers of the GM tetrahedra from the full head model were the inputs to the function. Center points outside of the original cropped GM volume were not assigned values and instead were set to MATLAB’s “not a number” (NaN) value. Because the interpolation uses a linear interpolation method, the values are slightly altered by surrounding values. This can result in shifting values towards a mean value, especially for the more extreme values surrounded by lower valued regions. Therefore, robust maximum of the electric field was not a valid measurement for the interpolated results. Instead, we calculated whole brain mean electric field strengths (|E|_mean_) in the comparison of the full vs. cropped head models. Nevertheless, the overall spread of the electric field is preserved during interpolation. An example of the original vs. interpolated simulation results can be seen in supplementary Figure S2.

## Results

### TES Muscle Results

We simulated 10 different TES montages for three muscle conditions (no muscle, isotropic muscle, and anisotropic muscle) to evaluate how muscle affects the estimate for the induced electric field. The simulation results across muscle conditions resulted in a similar spread of electric field, but with different intensities. An example for the montage using electrode pair 2–6 can be seen in Figure 2A. The areas of higher intensity were similar across all three conditions, but isotropic muscle had the highest electric field magnitude in those regions. The isotropic muscle condition had a maximum increase of 29.5% from the no muscle condition. On average, the maximum increase across all montages was 22.6% ± 5.2% (mean ± standard deviation). In contrast, the anisotropic muscle condition only had a maximum increase of 17.9% from the no muscle condition with a mean ± standard deviation of 13.5% ± 2.5% for all montages. These results can be seen in Figure 2B. Across all montages isotropic muscle > anisotropic muscle > no muscle for both the robust maximum of the electric field strength and whole brain mean electric field strength (Figure 2C).

**Figure 2.**
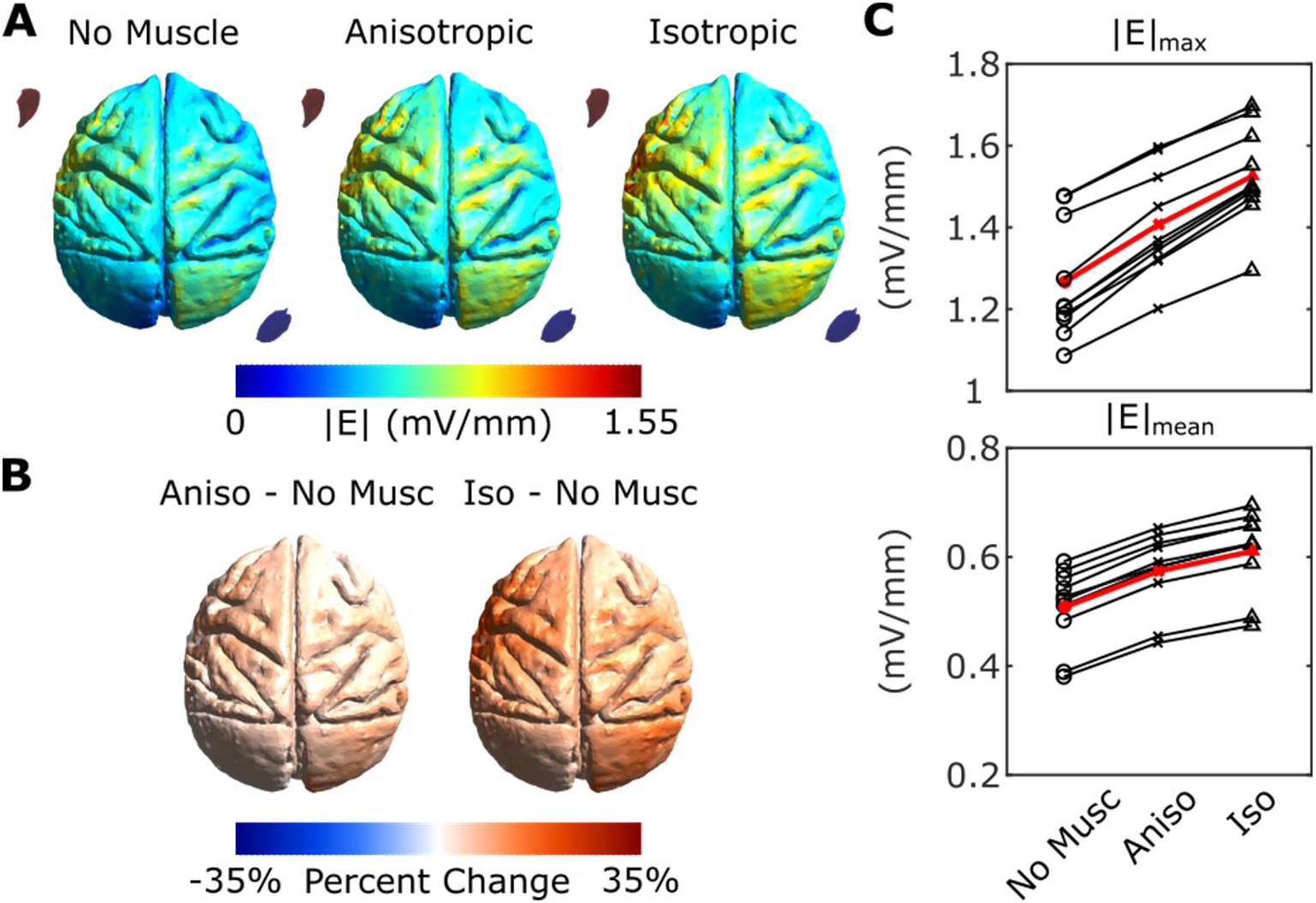
TES muscle condition simulation results. A) Example electric field magnitude (|E|) results from montage 2–6 for the three muscle conditions. The isotropic muscle resulted in areas of higher electric field strength than both the anisotropic muscle and no muscle conditions. B) The percent change of the difference between the models with muscle and the no muscle condition. There is a larger difference between the isotropic muscle condition and the no muscle condition (right) than between the anisotropic muscle condition and the no muscle condition (left). C) The maximum (top) and mean (bottom) electric field strength for each muscle condition. The results for each montage across conditions are connected by lines. And the averages for each condition across all montages are the red bold symbols and lines. The results show that for all 10 montages that electric field strength is highest for isotropic muscle and lowest no muscle for both maximum and whole brain mean.

### TMS Muscle Results

We ran TMS simulations for 36 coil positions and 8 coil orientations for the three muscle conditions (no muscle, isotropic muscle, and anisotropic muscle) to evaluate how muscle affects the estimate for the induced electric field. Three positions were excluded from the following results due to the coil intersecting the 3D head model (positions 1, 28, and 36). The simulation results across muscle conditions resulted in similar spread of electric field with similar intensities. An example using coil position 11 in the 0° orientation can be seen in Figure 3A. The difference in the electric field strength was negligible. The isotropic muscle condition had a maximum increase of 3.0% and maximum decrease of 6.0% from the no muscle condition. Similarly, the anisotropic muscle condition ranged from −6.1% to +3.3% change from the no muscle condition. These results can be seen in Figure 3B. Across all coil positions and orientations, there is almost no difference in the distribution of maximum electric field strength (Figure 3C).

**Figure 3.**
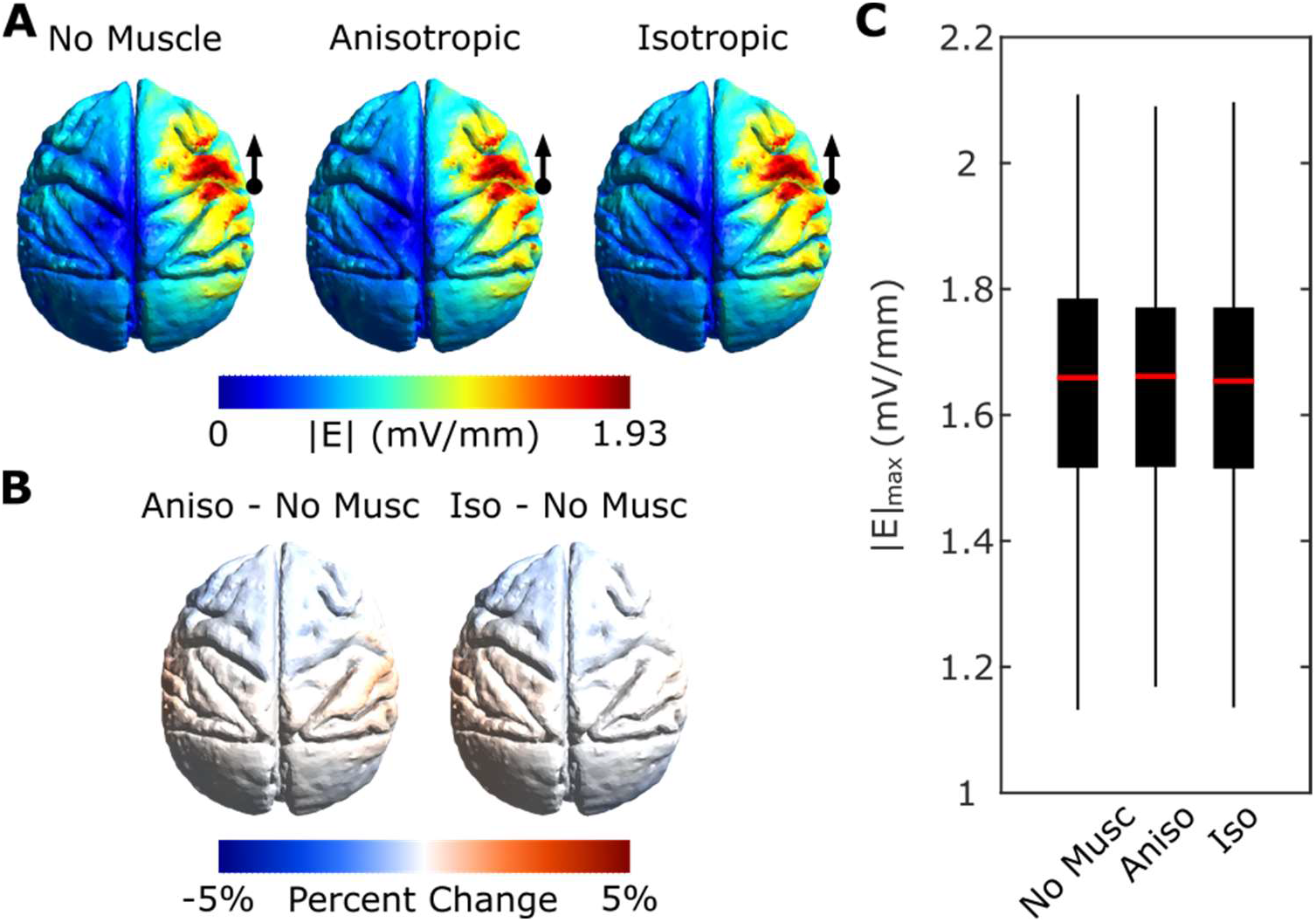
TMS muscle condition simulation results. A) Example electric field magnitude (|E|) results from position 11 in the 0° orientation (black dot and arrow indicate coil position and orientation) for the three muscle conditions. There is little noticeable difference in electric field strength and spread across the three muscle conditions. B) The percent change of the difference between the models with muscle and the no muscle condition. There is a negligible difference between the muscle conditions and the no muscle condition, although there are regions that are slightly higher or slightly lower than the no muscle condition. C) The electric field strength robust maximum for each muscle condition for all coil positions and orientations. The electric field strength shows little difference between muscle conditions even when all coil parameters are taken into account.

### TES Cropped Model Results

The same ten TES montages were run on the cropped head model to compare to the no muscle full head condition to understand how increasing the volume of the head model affects the simulation results. Once the initial simulations were completed, the cropped head model results were interpolated into the same GM elements present in the full head model to allow for a one-to-one comparison across elements. The simulated electric fields (Figure 4A) show higher intensity results in similar areas of the brain, however the cropped head model has higher electric field strength. This difference in electric field strength is clearly seen in Figure 4B where almost all elements show a higher electric field strength for the cropped model than for the full model. The electric field strength of the cropped head model compared to the full head model can be up to 13.5% ± 3.2% (maximum 16.9%, minimum 8.8%) higher relative to maximum of the cropped model. Additionally, the whole brain mean electric field strength is higher in the cropped model than in the full model, as seen in Figure 4C. We compared every tetrahedral element between the full and cropped models for all ten montages which are shown in supplementary Figure S3.

**Figure 4.**
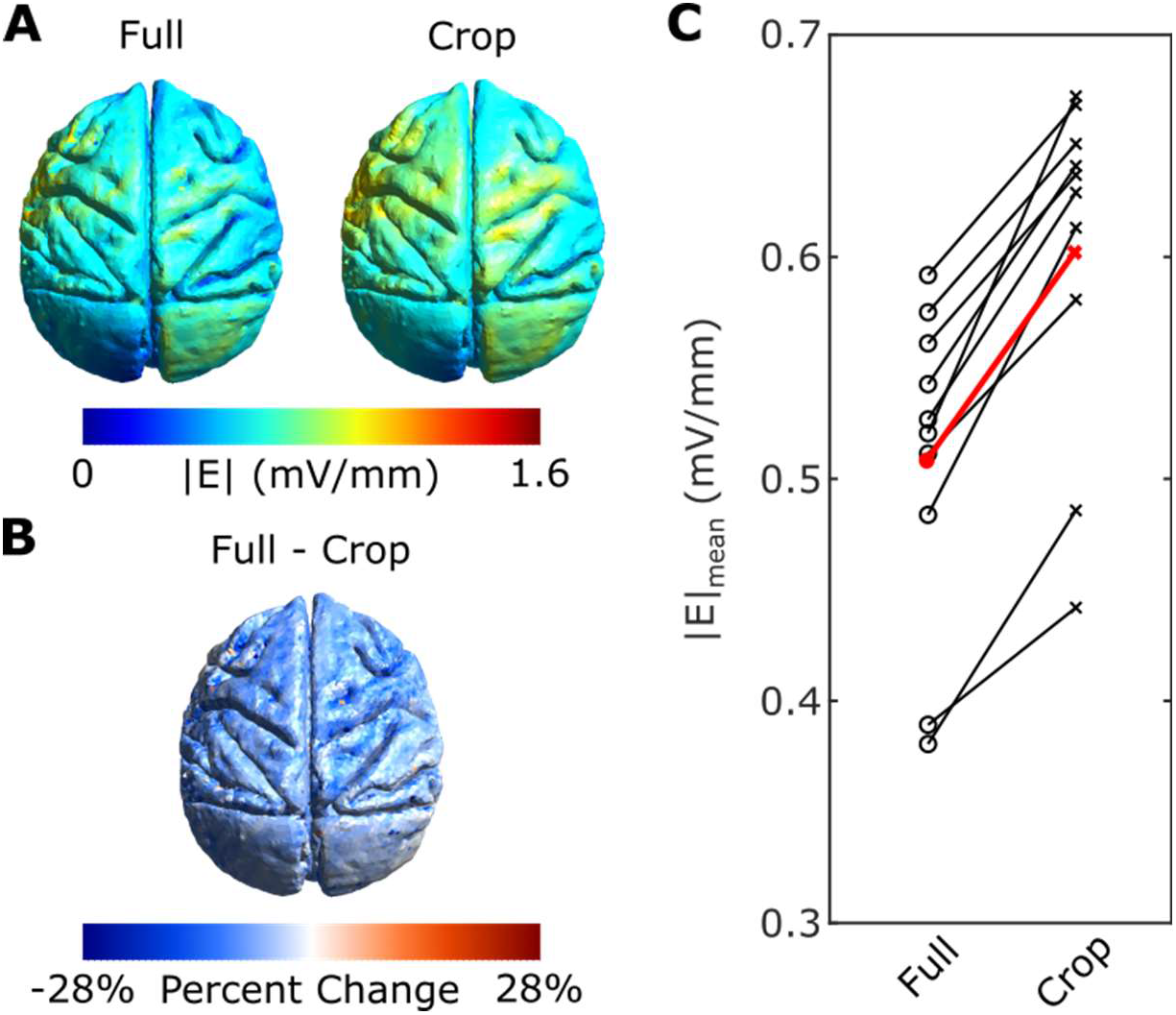
TES cropped vs. full simulation results. A) Example electric field magnitude (|E|) results from montage 2–6 for the interpolated cropped model and full model. The cropped model resulted in areas of higher electric field strength than the full model. B) The percent change of the difference between the models. The cropped model has consistently higher electric fields than the full model for most GM elements. C) The whole brain mean electric field strengths for full and cropped models. The results for each montage across conditions are connected by lines and show that the cropped model has a higher mean electric field strength than the full model in all cases. The bold red line is the average of all montages for each condition.

### TMS Cropped Model Results

All of the same TMS simulations (36 positions and 8 coil orientations) were run on the cropped model to evaluate the effect of head volume on the TMS simulation results. To be consistent with the full head model, the same three positions (1, 28, and 36) were excluded from the results. As with TES, the cropped head model results were interpolated into the same GM elements present in the full head model to allow for a one-to-one comparison of results across elements. In the case of TMS, there is a larger spread of high electric field strength than seen in the cropped head model (Figure 5A). This is more easily seen in the difference mesh in Figure 5B; the full head model has a higher electric field strength compared to the cropped head in the hemisphere under the coil position. The electric field strength of the full head model can be up to 21.5% ± 3.0% (maximum 34.5%, minimum 17.3%) higher relative to the maximum of the cropped model across all coil positions and orientations. But the hemisphere contralateral to the coil placement has the opposite effect; the electric field strength is higher for the cropped model. This pattern was consistent for all TMS coil locations. The region under the coil was higher for the full head model, but the region of the brain opposite the coil was higher for the cropped head model. Despite the duality of the difference results, the whole brain mean of electric field strength is higher for the full head model (Figure 5C). We explored this further by looking at current streamlines from the current density simulation results (see supplementary Figure S4 for an example). We found that for a given starting position, the current will often wrap around the brain more times in the cropped model due to the limited size of the volume conductor. However, in the full head model, the current has space to travel through and therefore does not necessarily return to the opposite side of the brain as often.

**Figure 5.**
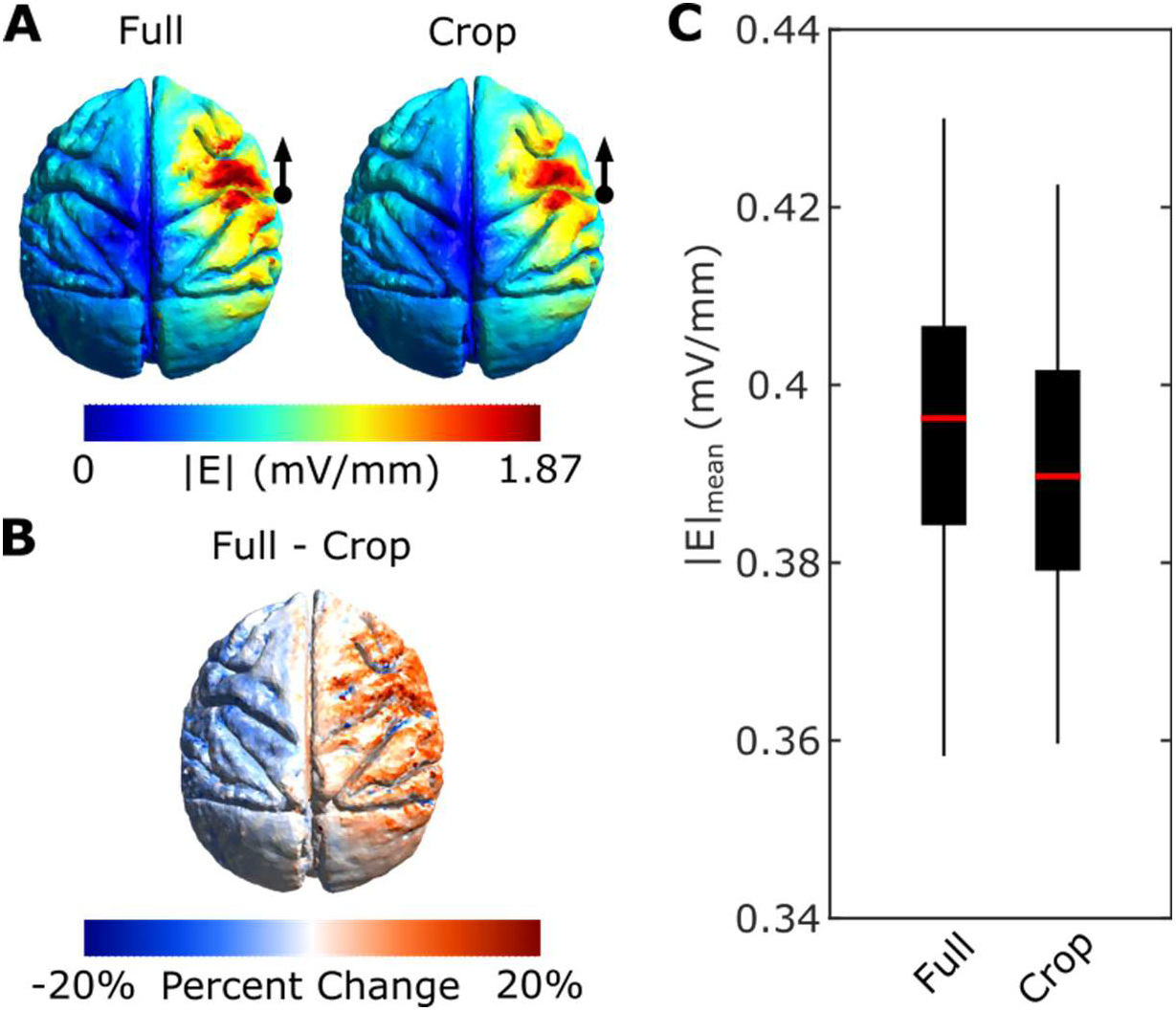
TMS cropped vs. full simulation results. A) Example electric field magnitude (|E|) results from position 11 in the 0° orientation (black dot and arrow indicate coil position and orientation) for the interpolated cropped model and full model. The cropped model has a similar distribution for electric field strength, but the magnitudes are different. B) The percent change of the difference between the models. The full model has consistently higher electric fields than the cropped model for the region nearest the coil, but lower electric fields for the opposite side of the brain. C) The whole brain mean electric field strength for full and cropped models. The results for each coil position and orientation show that the full model has a slightly higher mean electric field strength on average than the cropped model.

## Discussion

There are important implications in the accurate calculation of the electric field induced by NIBS for translational work between NHPs and humans (Alekseichuk et al., 2019; Johnson et al., 2020; Kar et al., 2017; Krause et al., 2017, 2019; Mueller et al., 2014; Opitz et al., 2016). Thus, it is crucial to evaluate how anatomical details in NHP head models affect electric field predictions. Namely, muscle tissue can significantly affect electric fields due to the large conductivity difference from skin, but has not been systematically evaluated yet. Furthermore, cropped head models are the standard in the field, but could potentially lead to misestimates of NIBS electric fields. Therefore, we systematically evaluated these two anatomical factors to determine what is required for accurate 3D NHP models. For this we developed a high resolution NHP head model from 10.5 T MR data with increased anatomical detail. We systematically evaluate the effect of both the inclusion of muscle anisotropy and head size on both TES and TMS electric fields.

In TES simulations, we respectively saw up to a 17.9% and 29.5% increase in the model with anisotropic and isotropic muscle compared to the model without any muscle. This difference occurs because the presence of muscle tissue subtracts from skin volume. Skin is highly conductive, thus a large portion of the electric current shunts through the skin (Vöröslakos et al., 2018). But, when a muscle is added into the model, it limits the skin shunting and increases current flow in the brain. In our study, we find that this effect is strongest for isotropic muscle. In the case of the anisotropic condition, the current travels more easily along muscle fibers. Since the fibers usually follow along the skull contour, rather than pointing inwards toward the skull, less current penetrates the skull than in the isotropic case. However, the skin-muscle conductivity mismatch is still present, so more current travels into the muscle volume and is enough to increase the current flow into the brain compared to no muscle. In this study we used a 5 : 1 ratio for parallel to radial anisotropic conductivity. Theoretically, we expect that if there is a large enough ratio, the parallel current shunting would overwhelm the current boost from the skin interface and the current reaching the brain (and therefore the electric field strength in the brain) would be lower than the no muscle case. However, we do not expect this to happen in the biologically plausible range.

There are some fundamental differences between TMS and TES which would explain the different results between these modalities. In TES a current is directly applied to the head, and thus muscle affects this current flow. For TMS, the magnetic field passes through head and is not affected by the presence of muscle tissue. TMS-induced currents are instead derived from the electric fields generated from the time varying magnetic field in the brain. Consequently, the electric field in the brain is mainly determined by the tissue boundary of CSF and GM (Miranda et al., 2003; Thielscher et al., 2011). Therefore, the electric field in the brain will be relatively unchanged in the different muscle models. This is verified by our results where the largest change is only −6.1%. The slight difference in results likely comes from the secondary electric field component due to charge accumulation at tissue interfaces. There are slight differences with the presence of muscle, but these effects can be ignored from a modeling perspective.

The comparison between cropped and full head models for TES simulations supported our prediction that a cropped head model will have higher electric fields. In our previous work we demonstrated this effect by comparing different species (Alekseichuk et al., 2019). Here we investigate this effect for an individual with differences in the completeness of the head model. The decrease in electric field strength occurs because the finite amount of current being injected into the head from the electrodes spread more in the full head due to the larger volume conductor. This generates a lower current density everywhere in the brain.

In TMS, our previous work indicated that a larger head could capture more energy from the coil (up to a point) leading to higher electric fields. This was generally supported by our results for average electric field strength. Interestingly we saw a pattern where the full head model has a higher electric field strength underneath the coil, but lower in other areas. Investigating the current streamlines, we found that while the larger volume in the full head captures more energy and explains the higher electric fields under the coil, the currents have more room to spread in areas away from the coil. This leads to lower electric fields in the contralateral hemisphere to the TMS coil. Conversely, the cropped model traps the energy it captures and the current looping concentrates energy in other regions of the brain. Overall, this pattern is not critical for translational work, because the regions where the cropped head electric fields are higher than the full head are only in areas of low electric field strength, where the brain is less affected by the stimulation.

## Conclusion

In summary, our work demonstrates that anatomical detail in NHP models affects the estimated NIBS induced electric fields. Muscle tissue is an important factor in TES simulations which could affect the estimated electric field strength in the brain up to 29.5%. Full head models affect the results for both TES and TMS. The larger volume results in opposing effects for TES and TMS on the electric field strength up to 13.5% and −21.5%, respectively. Our results can help experimental planning of NIBS research in NHPs as well as the translation of findings between NHPs and humans.

