## Supplementary Material for "Systematic evaluation of anatomical details on transcranial electric stimulation and transcranial magnetic stimulation induced electric fields in a non-human primate model"

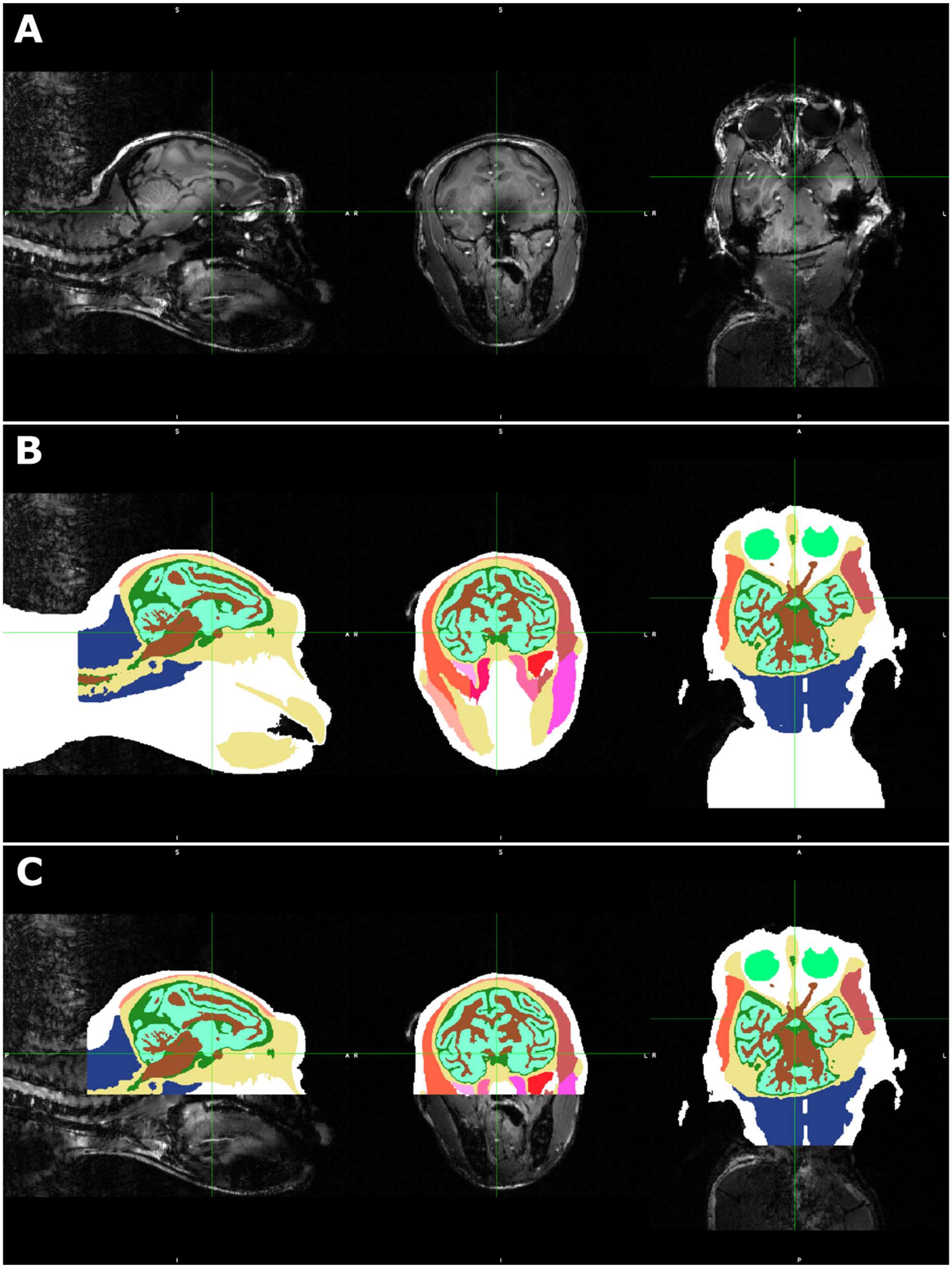


*Figure S1. Full and cropped masks created from 10.5 T MR image. A) 10.5 T MR image. B) The mask for the full head was created using a combination of tools and hand segmentation. From inside and moving outward the tissues are as follows: brown*

*is WM, light blue is GM, dark green is CSF, tan is skull, reds/pinks outside the skull are head muscles, dark blue is neck muscles, light green is eyes, white is skin. C) The cropped mask was made directly from the full mask. Two planes were selected and all mask labels beyond those planes were removed in MATLAB.*

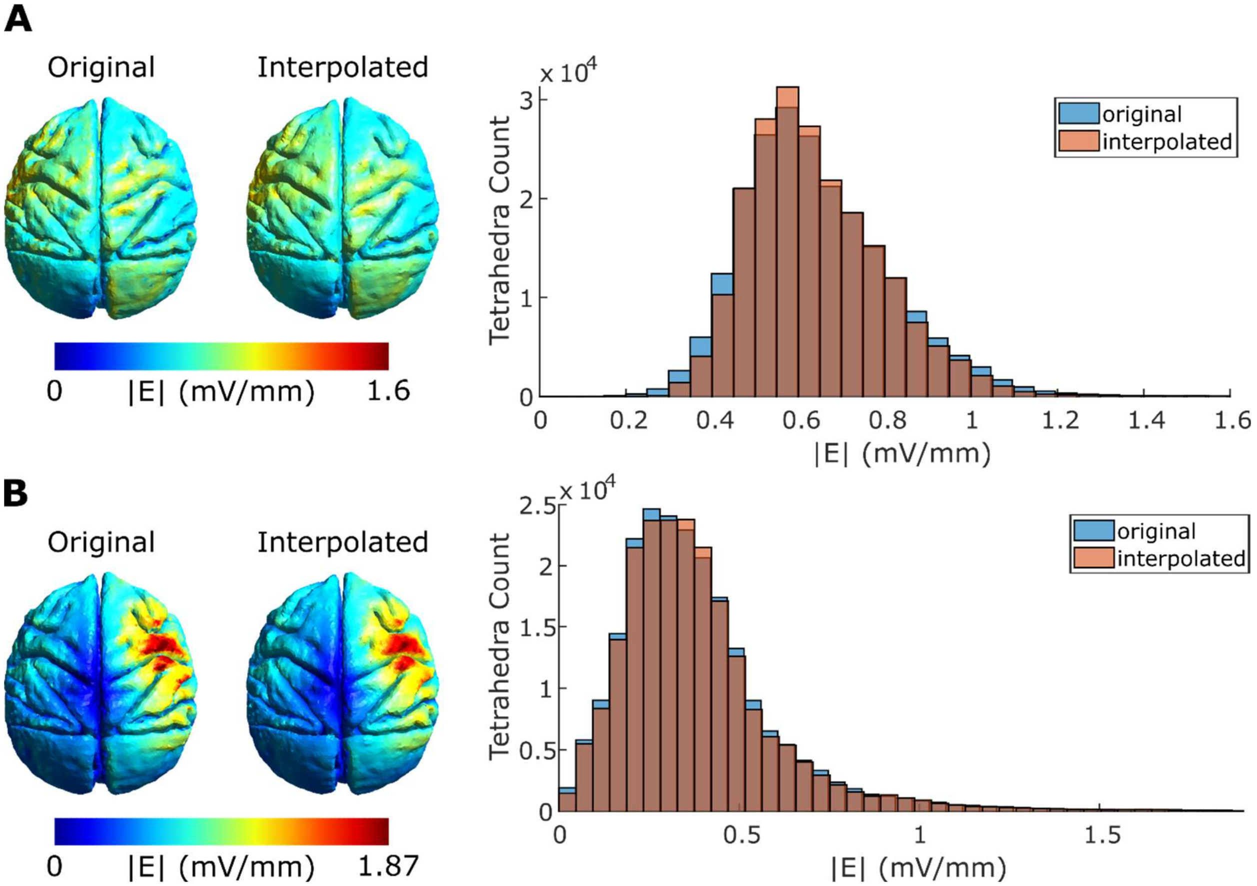


*Figure S2. Comparison of the original cropped head model and the interpolated model. The centers of the GM tetrahedra from the full head model were the coordinates input into the interpolation function. This enabled a one-to-one comparison of all tetrahedra between the full and cropped models. Overall, 6573 tetrahedra from the full model were outside of the original cropped GM mesh, and thus were not assigned a value (NaN in MATLAB). This accounts for 3.01% of all full model GM tetrahedra. A) An example TES simulation (montage 2–6). The electric field strength (|E|) distribution is very similar between the original cropped model and the interpolated model (left). The histograms verify the distribution of electric field strength is very similar for both models (right). There is a slight shift from the edges towards the apex of the histogram in the case of the interpolated histogram. This is not unexpected because a linear interpolation method can bring extremes closer to the average. B) An example TMS simulation (position 11 with $0^o$ orientation). Again, the electric field strength distribution is similar for both models (left) and the histograms verify the similarity for all tetrahedron (right).*

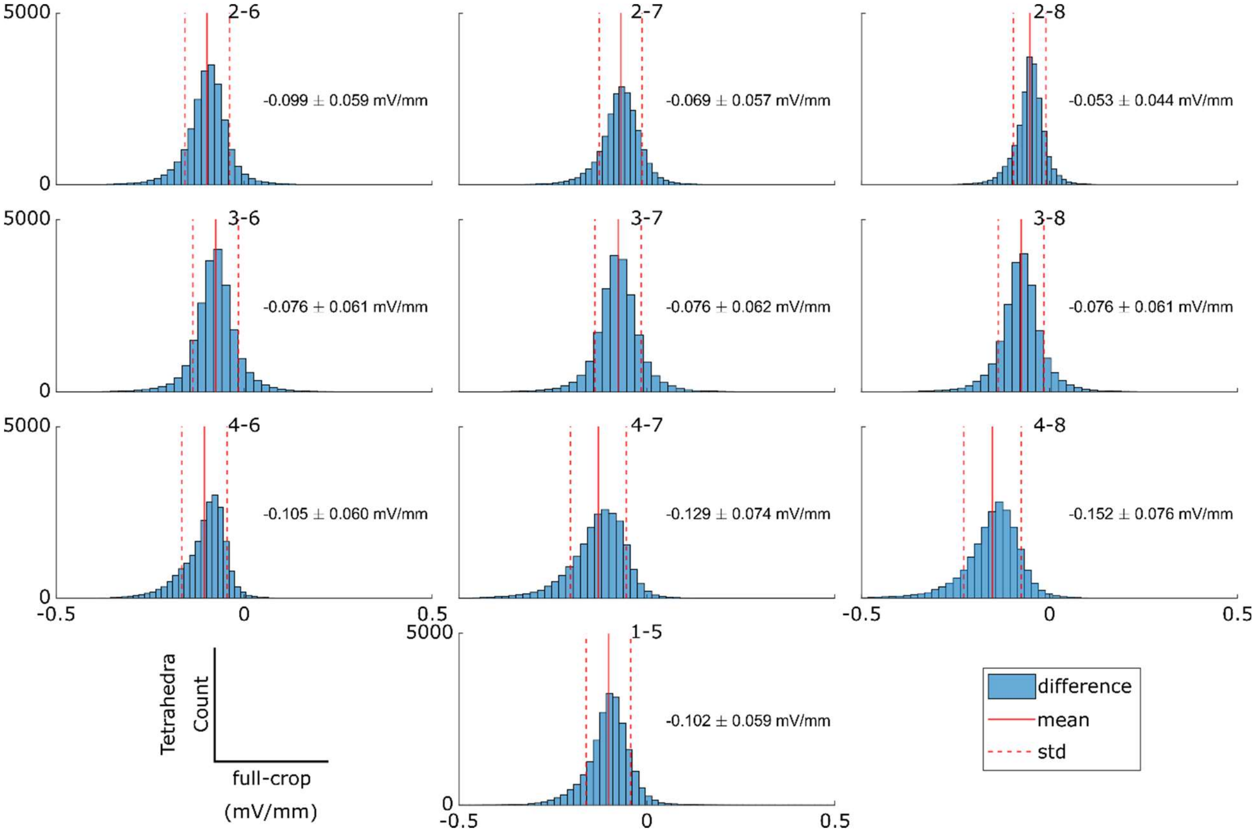


*Figure S3. Histograms of TES full – crop difference. The mean difference for each montage is indicated by the solid red line and the standard deviation is indicated by the dotted red line. Additionally, these values are explicitly stated as text on each plot. The histograms verify that the mean difference and mean + 1 standard deviation is less than 0. This confirms that the electric field strength for the majority of tetrahedra in the full model are less than the cropped model.*

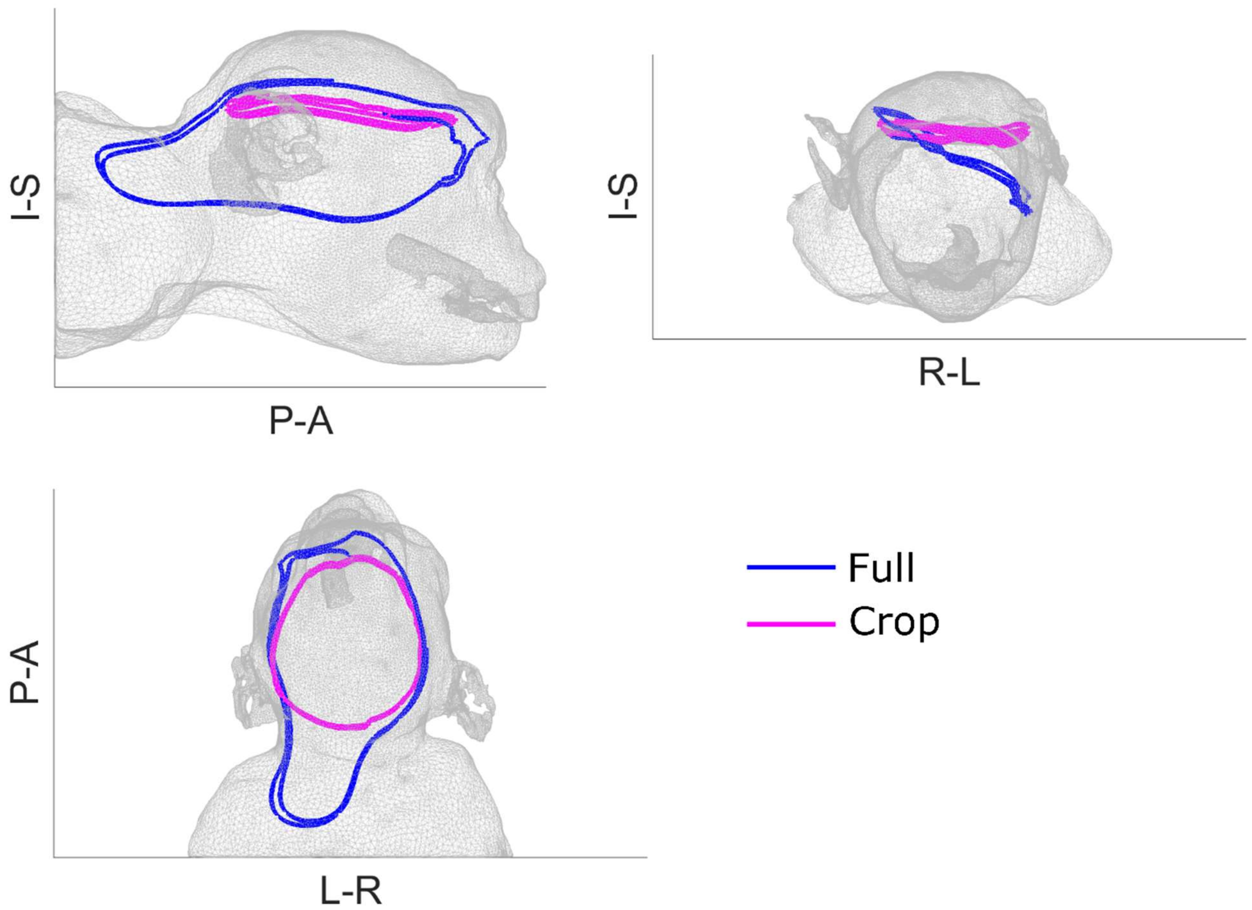


*Figure S4. Example current streamline comparison for full and cropped head models. These streamlines were generated with the MATLAB streamline function. The origin of the streamline is the same for both models, however they take dramatically different paths. In the full head (blue line), the current has a large volume that it is able to travel through. In contrast, the cropped head (pink line) shows that the current is trapped in a smaller volume, so it is forced to wrap around over and over in the same area. This could explain why the cropped model shows higher current density and therefore higher electric field strength values than the full head model in brain regions opposite the coil placement.*
